# Mitochondrial-nuclear interactions maintain geographic separation of deeply diverged mitochondrial lineages in the face of nuclear gene flow

**DOI:** 10.1101/095596

**Authors:** Hernán E. Morales, Alexandra Pavlova, Nevil Amos, Richard Major, Andrzej Kilian, Chris Greening, Paul Sunnucks

## Abstract

Metabolic processes in eukaryotic cells depend on interactions between mitochondrial and nuclear gene products (mitonuclear interactions). These interactions could play a direct role in population divergence. We studied the evolution of mitonuclear interactions in a widespread passerine that experienced population divergence followed by bi-directional mitochondrial introgression into different nuclear backgrounds. Using >60,000 SNPs, we quantified patterns of nuclear genetic differentiation between populations that occupy different climates and harbour deeply divergent mitolineages despite ongoing nuclear gene flow. Analyses were performed independently for two sampling transects intersecting mitochondrial divergence in different nuclear backgrounds. In both transects, low genome-wide nuclear differentiation was accompanied by strong differentiation at a ~15.4 Mb region of chromosome 1A. This region is enriched for genes performing mitochondrial functions. Molecular signatures of selective sweeps in this region alongside those in the mitochondrial genome suggest a history of adaptive mitonuclear co-introgression. The chromosome 1A region has elevated linkage disequilibrium, suggesting that selection on genomic architecture may favour low recombination among nuclear-encoded genes with mitochondrial functions. In this system, mitonuclear interactions appear to maintain the geographic separation of two mitolineages in the face of nuclear gene flow, supporting mitonuclear co-evolution as an important vehicle for climatic adaptation and population divergence.

## Introduction

Genomic studies of early stages of population divergence enhance our understanding of the genetic basis of local adaptation, reproductive isolation and speciation (Harrison & Larson 2016; Payseur & Rieseberg 2016; Seehausen *et al.* 2014). Genomic differentiation between closely related populations is often heterogeneous: low differentiation across most of the genome is accompanied by high differentiation at barrier loci, i.e. genes involved in local adaptation and/or reproductive isolation (Wu 2001). Highly differentiated genes are sometimes packed into distinctive chromosomal regions, commonly referred to as genomic islands of differentiation (Ravinet *et al.* 2017; Wolf & Ellegren 2016). Genomic islands of differentiation can evolve during divergence-with-gene-flow in order to prevent recombination between barrier loci and reduce gene flow for them (Nosil *et al.* 2009), or during periods of population isolation as a byproduct of ancient directional or background selection that resulted in low genetic variation within genomic regions (Cruickshank & Hahn 2014; Nachman & Payseur 2012; Noor & Bennett 2009). To date, efforts to uncover the genomic basis of differentiation in the wild have mostly focused on nuclear genes (e.g. Jones *et al.* 2012; Marques *et al.* 2016; Soria-Carrasco *et al.* 2014). In contrast, few studies have approached the co-evolution of nuclear and mitochondrial genomes during the early stages of divergence in natural populations (e.g. Bar-Yaacov *et al.* 2015; Baris *et al.* 2017; Boratyński *et al.* 2016; Gagnaire *et al.* 2012; Sambatti *et al.* 2008) and none have tested whether genomic islands of differentiation are enriched for nuclear genes with mitochondrial functions. Genes of the mitochondrial genome (mtDNA) and nuclear-encoded genes with mitochondrial functions (N-mt genes) are nevertheless prime candidates to drive population divergence because they are responsible for maintaining essential functions in energetics, metabolism, and gene regulation (Allen 2003; Horan *et al.* 2013). Most significantly, mtDNA and N-mt genes co-encode and regulate the oxidative phosphorylation (OXPHOS) complexes, which serve as the primary source of energy availability in the cell (Bar-Yaacov *et al.* 2012). Accordingly, genetic variation in mtDNA has strong fitness effects, often expressed through mitonuclear interactions (Ballard & Pichaud 2014; Wolff *et al.* 2014). The interactions and resulting fitness effects can be environment-dependent (Arnqvist *et al.* 2010; Hoekstra *et al.* 2013).

Mitonuclear interactions occur despite mitochondrial and nuclear genomes having different modes of inheritance, recombination and mutation rates, implying that their co-evolution is enforced by natural selection (Dowling *et al.* 2008; Rand *et al.* 2004). Selection can favour the formation of mitonuclear interactions during population divergence in three main ways. First, selection to maintain metabolic functionality drives co-evolution of mtDNA and N-mt genes: in response to rapid accumulation of slightly deleterious mutations and/or adaptive variation in mtDNA, N-mt genes evolve compensatory or epistatic changes (James *et al.* 2016; Osada & Akashi 2012; Sloan *et al.* 2016). Thus, mitonuclear co-evolution can result in populations having different co-adapted sets of mtDNA and N-mt alleles. Second, environmental variation can drive selection for locally-adapted metabolic phenotypes, modulated by mitonuclear interactions (Burton *et al.* 2013; Hill 2015). For example, variation in mitochondrial coupling between substrate oxidation and production of energy (ATP synthesis) can be adaptive in environments that differ in their thermal profile and nutrient availability (Das 2006; Lowell & Spiegelman 2000; Stier *et al.* 2014). Thus, environmental variation can act as an extrinsic barrier to gene flow, and result in diverging populations having different co-adapted sets of mtDNA and N-mt alleles. Third, when two populations with different sets of co-adapted mitonuclear alleles admix (e.g. after secondary contact or during mitochondrial introgression or isolation-with-migration), poor fitness of hybrid mitonuclear genotypes can drive selection against mitonuclear incompatibilities (Burton & Barreto 2012). To reduce fitness-loss through the production of hybrids with incompatible mitonuclear combinations, mechanisms could evolve to decrease mating between incompatible lineages (Burton & Barreto 2012; Burton *et al.* 2013), and/or to suppress recombination between co-adapted N-mt alleles (Lindtke & Buerkle 2015; Ortiz-Barrientos *et al.* 2016). Thus, selection against genetic incompatibilities and/or for recombination-inhibiting genomic architecture may act as intrinsic barriers to gene flow, leading to populations with different combinations of mtDNA and N-mt alleles.

The endemic eastern Australian songbird *Eopsaltria australis* (Eastern Yellow Robin; hereafter EYR) provides an excellent model in which to study mitonuclear interactions during divergence with gene flow, because it has two diverged populations that experienced extensive mitochondrial introgression from each other (Morales *et al.* 2017b; Pavlova *et al.* 2013; Fig. S1). Coalescent modelling indicates that northern and southern populations split approximately 2 million years ago, causing divergence in mtDNA lineages (mitolineages; now 6.8% divergent) and in nuclear DNA (nDNA; mean genome-wide G_ST_ = 0.084) (Morales *et al.* 2017b). Subsequently, ~270 thousand years ago, the northern mitolineage (mito-A) introgressed southwards through the relatively arid and climatically variable inland range, and more recently ~90 thousand years ago, the southern mitolineage (mito-B) introgressed northwards through the more temperate and climatically stable coastal range. This history approximates a ‘natural experiment’, in which a pattern of mitochondrial divergence between lineages occupying contrasting climates occurs in two different nuclear backgrounds (Fig. S1). Following southwards mito-A introgression, ~60 thousand years ago the southern population began to diverge into an inland (more arid) mito-A-bearing sub-population and a coastal (more temperate) mito-B-bearing sub-population. These two subpopulations are connected by male-mediated gene flow (Morales *et al.* 2017b). Signatures of positive selection on six amino acids of mitochondrially-encoded OXPHOS genes support non-neutral mtDNA divergence between the mitolineages (Lamb *et al.* submitted; Morales *et al.* 2015). Moreover, signatures of selective sweeps in the mitogenome and significant correlations of mitolineage distributions with climatic variation after controlling for the effects of geography suggest that one or both mitochondrial introgression events may have been adaptive (Morales *et al.* 2015; Pavlova *et al.* 2013). We hypothesize that non-neutral inland-coastal mitolineage differentiation in the face of nuclear gene flow should be accompanied by corresponding differentiation in regions of the nuclear genome linked to mitochondrial function, reflecting selection for mitonuclear interactions. Specifically, if the initial north-south population divergence generated N-mt genes co-adapted to their regional mitochondrial type, mitochondrial introgression should have been accompanied (or soon followed) by introgression of co-evolved alleles of N-mt genes (i.e. mitonuclear co-introgression; e.g. Beck *et al.* 2015; Boratyński *et al.* 2016).

Here we examined these predictions using genomic data collected along two geographic transects intersecting the EYR’s inland-coastal mitochondrial divergence in both northern and southern nuclear backgrounds. We used 60,444 SNPs to detect loci strongly differentiated between inland and coastal populations defined by their mitolineage (i.e. outlier loci) in each transect/nuclear background. We quantified clinal variation among a subset of 2,494 loci to approximate genome-wide and locus-specific patterns of nuclear genetic introgression. We also quantified the level of nuclear admixture in each transect for neutral and non-neutral loci. Moreover, using the zebra finch genome as a reference, we investigated if outlier loci are arranged into genomic islands of differentiation, and tested if these islands are enriched with N-mt genes, as expected if mitonuclear interactions drive population divergence. Furthermore, we tested if genomic islands of differentiation have low genetic diversity and high linkage disequilibrium, as expected under mitonuclear co-introgression generating nuclear selective sweeps paralleling those previously observed in the mitogenomes (Morales *et al.* 2015; Morales *et al.* 2017b). Finally, to test whether the N-mt OXPHOS genes located within the islands participate in protein-protein interactions with the mitochondrially-encoded subunits under positive selection (Lamb *et al.* submitted; Morales *et al.* 2015), we mapped OXPHOS protein sequences of the reference genome and EYR mitochondrial OXPHOS protein sequences under selection, onto 3D proteinstructure models of OXPHOS complex I.

## Results

### Narrow mitolineage contact zones at regions of environmental divergence occur in both nuclear backgrounds

A total of 407 EYR individuals sampled across the species’ range were categorized for mitochondrial lineage membership by ND2 mitochondrial DNA sequencing (262 inland mito-A and 145 coastal mito-B, coloured red and blue respectively on Fig. 1A; Table S1). Based on inferences of nuclear background from earlier analyses (Morales *et al.* 2017b), sampling efforts were focussed on two transects ~700 km apart, one in the northern nuclear background and one in the southern (squares and circles, respectively on Fig. 1A). In each transect, a narrow (~ 20 – 40 km) contact zone between mitolineages, defined based on presence of both mitolineages in a single sampling region (Fig. S12-S14), is located in a region of climatic transition (Fig. 1A). The correlation between mitolineage distribution and climatic variation is stronger in the southern than northern transect (Fig. 1B).

**Figure 1.**
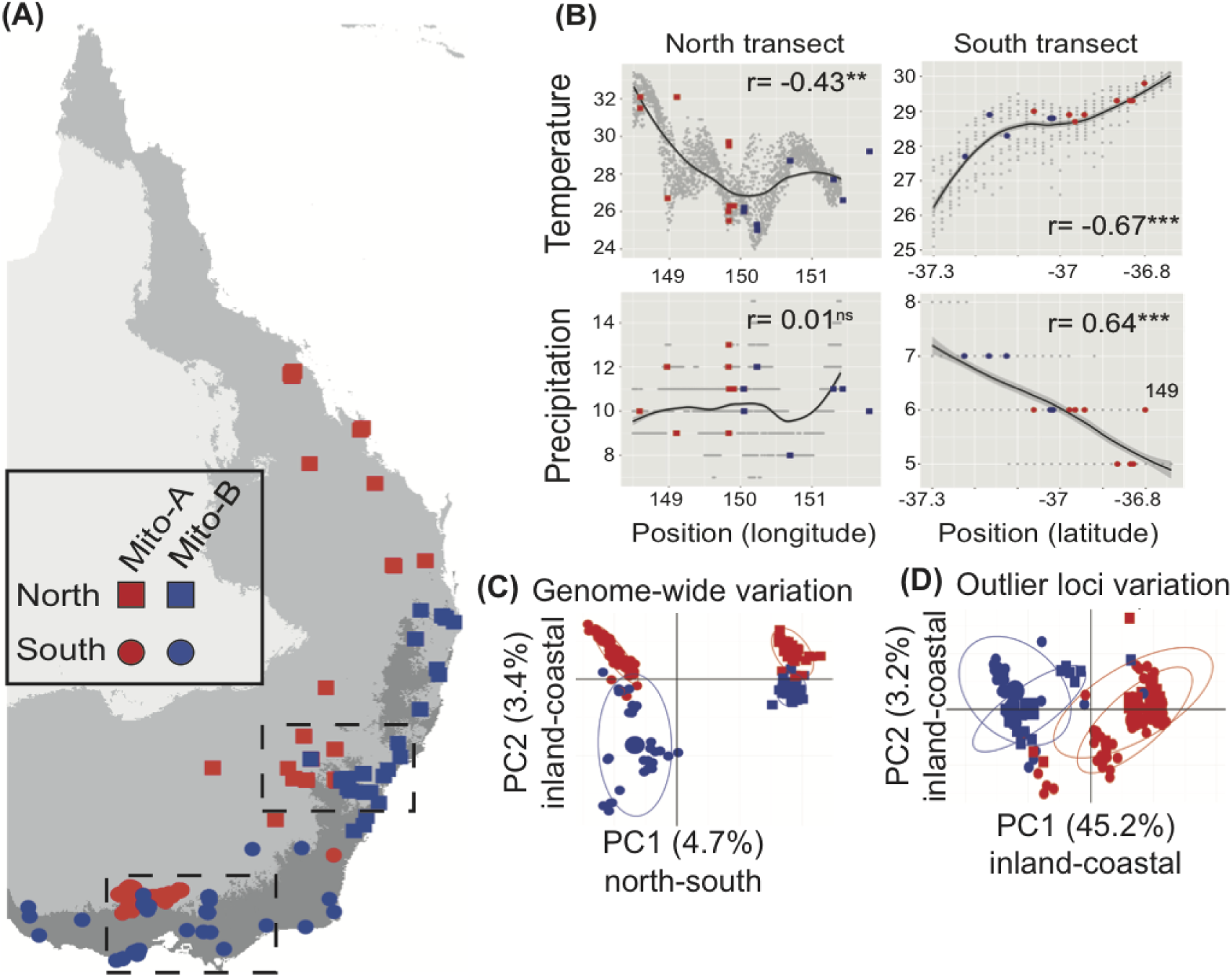
Geographic distribution of mitochondrial and nuclear genetic variation across the Eastern Yellow Robin (EYR) range. **(A)** Geographic distribution of EYR 6.8% divergent mitolineages, inland mito-A (red) and coastal mito-B (blue), in different nuclear backgrounds, north (squares) and south (circles; same symbols apply to the remaining panels). Sampling was focused along two transects (black dashed rectangles), one in each nuclear background. Shades of grey represent maximum temperature of warmest month (BIOCLIM 5): dark-grey < 28C°, medium-grey 28-33C° and light-grey > 33 C°. The two mitolineages meet in a narrow contact zone of climatic divergence over 1,500 km. **(B)** We estimated the Pearson’s correlation coefficient between mitolineage distribution and climatic variation along each transect (three asterisks indicate P < 0.001, two asterisks- P < 0.01, ns- P > 0.05). Y-axes show climatic variables: maximum temperature of warmest month in °C (BIOCLIM 5) and precipitation of the driest month in millimetres (BIOCLIM 14). X-axes geographic position (latitude or longitude) in degrees. Grey points represent background climatic variation of random sampled points in each transect. Black lines represent average background climatic variation obtained by fitting a linear model of background points against position **(C)** Principal component analysis (PCA) of nuclear genome-wide variation (60,444 SNPs) of all EYR samples showing two major axes of genetic differentiation, corresponding to north-south (along PC1) and inland-coastal (along PC2) directions (PC1 vs PC2 plot built using only non-outlier loci depict the same structure, Fig. S3). **(D)** PCA of 565 SNP loci that are outliers between populations harbouring different mitolineages showing two major axes of genetic differentiation in an inland-coastal direction. Notably, outlier loci contain no signal of the north-south genome-wide divergence along the two first PCA axes. Ellipses in both PCA plots capture 80% of the data spread for each group.

### Strong genetic differentiation of outlier loci suggests a history of mitochondrial-nuclear co-introgression

We obtained 60,444 SNPs by performing complexity-reducing representative sequencing of the genome (DArTseq; Kilian *et al.* 2012) for 164 individuals (100 mito-A individuals and 64 mito-B). Most individuals were located along the two transects (50 in the northern, 103 southern). We used three methods differing in approaches and assumptions to identify outlier loci between different mitolineage-bearing populations, independently for each transect (see Methods). All three methods take into account potential confounding factors (e.g. genetic drift) to distinguish true outliers from false positives (Hoban *et al.* 2016). We first calculated *F*_ST_-values comprising individuals sampled within 40 km of the contact zone between different mitolineage-bearing populations; we considered loci in the top 1% of the distribution to be outliers. Genome-wide genetic differentiation between mito-A- and mito-B-bearing populations was lower in the northern transect (all loci: mean *F*_ST_ = 0.02, sd = 0.08, 1% upper quantile = 0.43) than southern (mean *F*_ST_ = 0.05, sd = 0.01, 1% upper quantile = 0.51) (Fig. S2). Then, we used two methods specifically designed for outlier discovery, BayeScEnv (Villemereuil & Gaggiotti 2015) and PCAdapt (Duforet-Frebourg *et al.* 2016). Each of the three methods identified similar numbers of outliers in each of the transects, with appreciable commonality of the loci identified among methods and in both transects (Table 1). Genetic differentiation for outlier loci was more than 12 times greater than the average genome-wide differentiation (mean outlier *F*_ST_ = 0.44 in the north, 0.56 in the south). Despite strong differentiation, no loci were completely fixed between different mitolineage-bearing populations in a transect.

**Table 1.** **Number of outlier loci identified in northern and southern transects** using three different methods: *F*_ST_ = top 1% quantile of per-marker *F*_ST_ between different mitolineage-bearing populations within 40 km radius of the mitolineage contact zone centre; BayeScEnv = population differentiation with mitolineage membership used as a binomial environmental variable; and PCAdapt = population differentiation across the first latent factor (K = 1; akin to the first PCA axis, see Methods), which capture inland-coastal differentiation.

| Method | North | South | North/South in common |
| --- | --- | --- | --- |
| $F_{ST}$ | 315 | 285 | 210 |
| BayeScEnv | 227 | 303 | 191 |
| PCAdapt | 323 | 235 | 171 |
| Total | 443 | 357 | 565 |
| In common | 185 | 203 | 155 |

A principal component analysis (PCA) of genome-wide variation (i.e. all SNPs) for all 164 samples revealed two major axes of EYR genetic differentiation: north-south (PC1 = 4.7%) and inland-coastal (PC2 = 3.4%; Fig. 1C). The inland-coastal differentiation was observed not only in the southern nuclear background, as in an earlier study using a smaller set of nuclear markers (Morales et al., 2017b), but also for the northern nuclear background. This differentiation is consistent with two episodes of mitochondrial introgression causing inland-coastal nuclear divergence in both northern and southern populations (Fig. S1). The same structure was detected by the first two axes of PCA of non-outlier loci, presumably reflecting neutral processes (Fig. S3). In marked contrast to the genome-wide PCA, a PCA of outlier loci (565 SNPs identified in common between transects) shows that genetic variation along the first two PC axis for these loci is exclusively structured in the inland-coastal direction with no northsouth variation observed at all (PC1 45.2%, PC2 3.2%; Fig. 1D). This pattern in outlier loci strongly resembles that of the mitochondrial genome where genetic variation within each mitolineage is extremely low across the entire species range (Morales *et al.* 2015). The existence of this subset of nuclear genes that correlate so strongly with mitochondrial variation supports the possibility that some of the nuclear outlier loci co-introgressed with mtDNA (Fig. S1). The pattern of mitonuclear co-introgression is also supported by the strong correlation of the frequencies of the same outlier alleles between nuclear backgrounds (north vs. south mito-A-bearing populations: Pearson’s r=0.98 and 0.99 for mito-B, significantly (P < 0.001) higher than correlations for random nonoutlier loci: 0.74 and 0.78 respectively, Fig. S4).

### Highly differentiated loci are concentrated in two genomic islands of differentiation

We mapped the SNP-containing loci to the reference genome of the zebra finch, *Taeniopygia guttata* (Warren *et al.* 2010), obtaining unique, high-quality hits for 35,030 loci. The loci that were strongly differentiated between different mitolineage-bearing populations appeared clustered into certain genomic locations, in both transects (Fig. 2; Fig. S5-S7). We tested for the presence of statistically-supported genomic islands of differentiation between different mitolineage-bearing populations in each transect using a Hidden Markov Model (HMM), and identified two significant genomic islands of differentiation: one on chromosome 1A (~15.4 Mb long; approximate zebra finch genomic coordinates: 44,200,000 - 59,600,000) and one on chromosome Z (~0.75 Mb long; 2,000,000 - 2,750,000) (Fig. 2).

**Figure 2.**
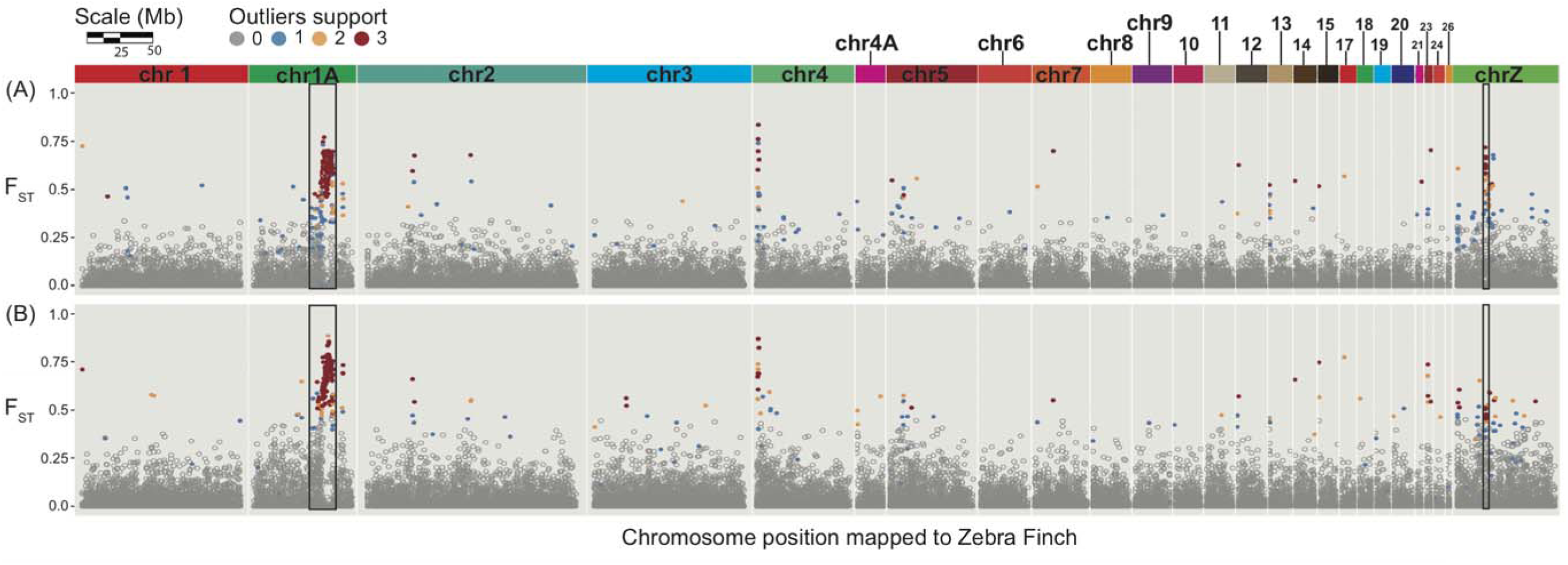
Heterogeneous genomic differentiation between different mitolineage-bearing populations for (A) northern and (B) southern transects. The Manhattan plots show *F*_ST_ on the y-axis and genomic position relative to the zebra finch reference genome on the x-axis for each SNP. The number of methods that support each outlier detected is indicated with different colours. Outlier detection methods employed were: *F*_ST_ outliers at fine spatial scales, BayeScEnv and PCAdapt (see Results for significance thresholds and Table 1 for number of outliers). Genomic regions with a significant cluster of contiguous genetic differentiation representing genomic islands of differentiation on chromosome Z and chromosome 1A were identified with Hidden Markov Models (black rectangles).

### The genomic island of differentiation on chromosome 1A has an overrepresentation of nuclear-encoded genes with mitochondrial functions

We tested if the two genomic islands of differentiation were significantly enriched for N-mt genes, in particular those encoding supernumerary subunits and assembly factors for OXPHOS complexes. The chromosome 1A island had a large and significant excess of N-mt genes: 32 genes compared to an average of 12.2 genes (sd = 5.6) in 3500 random regions of the same size across the genome (P < 0.001; Fig. S8; Table S3-S5). These 32 genes included a significant overrepresentation of N-mt genes that belong to the OXPHOS complexes: four genes compared to an average of 1.3 genes (sd = 1.2) in 3500 random genomic regions of the same size across the genome (P < 0.01; Fig. S8; Table S3-S5). Three of these genes (NDUFA6, NDUFA12, NDUFB2) encode supernumerary subunits required for OXPHOS complex I function (NADH-coenzyme Q oxidoreductase) and the fourth encodes the assembly chaperone FMC1 in OXPHOS complex V (F_1_F_o_-ATPase).

Of the remaining 28 N-mt genes in the island of differentiation, three more (in common with the OXPHOS ones) are directly involved in cellular respiration, eight have regulatory functions including mtDNA transcription, translation and replication, and the remaining 17 perform other mitochondrial functions. One of the nuclear-encoded regulatory genes located in this island, YARS2 (mitochondrial tyrosyl- tRNA synthetase), interacts with the mitochondrially-encoded tRNA-Tyrosyl, previously shown to have fixed differences between EYR mitolineages (Morales *et al.* 2015). In humans, a pathogenic mutation in YARS2 has been implicated multiple diseases caused by mitochondrial dysfunction (Riley *et al.* 2010). Moreover, controlled experiments have shown that *Drosophila* hybrids harbouring mitonuclear incompatible combinations of these two tRNA genes suffer delayed development and reduced fecundity, exacerbated by high temperatures (Hoekstra *et al.* 2013; Meiklejohn *et al.* 2013). In contrast to the island of differentiation on chromosome 1A, that on the Z chromosome contained only one N-mt gene (SLC25A46, which promotes mitochondrial fission) and no OXPHOS genes.

We used a recently-published 3D protein-structure model of OXPHOS complex I (Zhu *et al.* 2016) (Fig. S9) to infer the location in OXPHOS complex I of the three nuclear chromosome 1A-encoded subunits located within the island of divergence, as well as the three mitochondrially-encoded subunits previously inferred to be under positive selection in EYR (ND4, ND4L and ND5; Lamb *et al.* submitted; Morales *et al.* 2015). The three complex I OXPHOS subunits encoded within the chromosome 1A island (Table S5) occur at regions critical for energy transduction: NDUFA6 has an ancient role in stabilising the main interface between the hydrophobic and hydrophilic regions in complex I, NDUFB2 interacts directly with the mitochondrially-encoded ion pump ND5, and NDUFA12 binds a phosphopantetheine-containing ACP subunit (SDAP-α) implicated in regulating and stabilising the complex (Angerer *et al.* 2014; Fiedorczuk *et al.* 2016; Ostergaard *et al.* 2011; Yip *et al.* 2011; Zhu *et al.* 2016). The positions of these subunits in the protein complex suggest that, while they may not make direct protein-protein interactions with the three mitochondrially-encoded subunits under positive selection, they may alter functioning of the complex through driving long-range conformational changes dependent on the nature of the mitochondrial-encoded subunits (Fig. S9).

### Restricted introgression between mitolineages of loci within the chromosome 1A genomic island of differentiation

We used geographic cline analysis to test whether levels of nuclear introgression between different mitolineage-bearing populations varied across loci in each transect. Our expectation was that some nuclear gene regions with strong evolutionary interactions with mitochondrial variation under selection should show clinal variation across transects similar to that seen in mtDNA. We estimated variation in allelic frequencies as a function of geographic distance across each transect (the northern one ran 307 km east-west, the southern 174 km north-south) with a subset of the mapped SNPs (2,494 loci, including 42 outliers, one sampled randomly every 100 Kb, plus mtDNA) and compared its fit to clinal and non-clinal models. Within this subsample, we identified a total of 56 significantly clinal loci in the north and 146 in the south (Fig. 3A). Proportionately fewer non-outlier loci were clinal (1.3% of those tested in the north, and 4.3% in the south), in strong contrast to the very high proportion of outliers that were clinal (55% and 90% respectively). Clinal loci were significantly overrepresented in the chromosome 1A genomic island of differentiation, as expected if this genome region experiences particularly low introgression between different mitolineage-bearing populations (north: N = 14, t_27_ = -17.8, P < 0.001; south: N = 31, t_27_ = -19.3, P < 0.001). Not only were outliers disproportionately likely to be clinal, but their allelic frequency changes were much more geographically abrupt than those of clinal nonoutlier loci: the mean cline width of outlier loci was 42 km in the north and 64 km in the south, compared to 196 km and 184 km for non-outliers, respectively. However, the majority of clinal loci had confidence interval widths considerably greater than those of the mitochondrial clines (mtDNA cline widths: north = 5.9 km (2.1-20.5) south = 57.8 km (38.2-96.4)), indicating generally higher introgression for nDNA than mtDNA. Nonetheless, the presence of some overlap between nuclear and mitochondrial cline centres and widths signalled a subset of nuclear markers experiencing restricted introgression across the contact zones between different mitolineage-bearing populations (Fig. 3A). Loci in the chromosome 1A genomic island of differentiation showed the greatest overlap of cline parameters between mitochondrial and nuclear clines. Overall, our results suggest that, despite extensive genome-wide nuclear introgression between different mitolineage-bearing populations in each transect, a subset of outlier nuclear loci, including a disproportionate number in the chromosome 1A genomic island of differentiation, experience restricted introgression.

**Figure 3.**
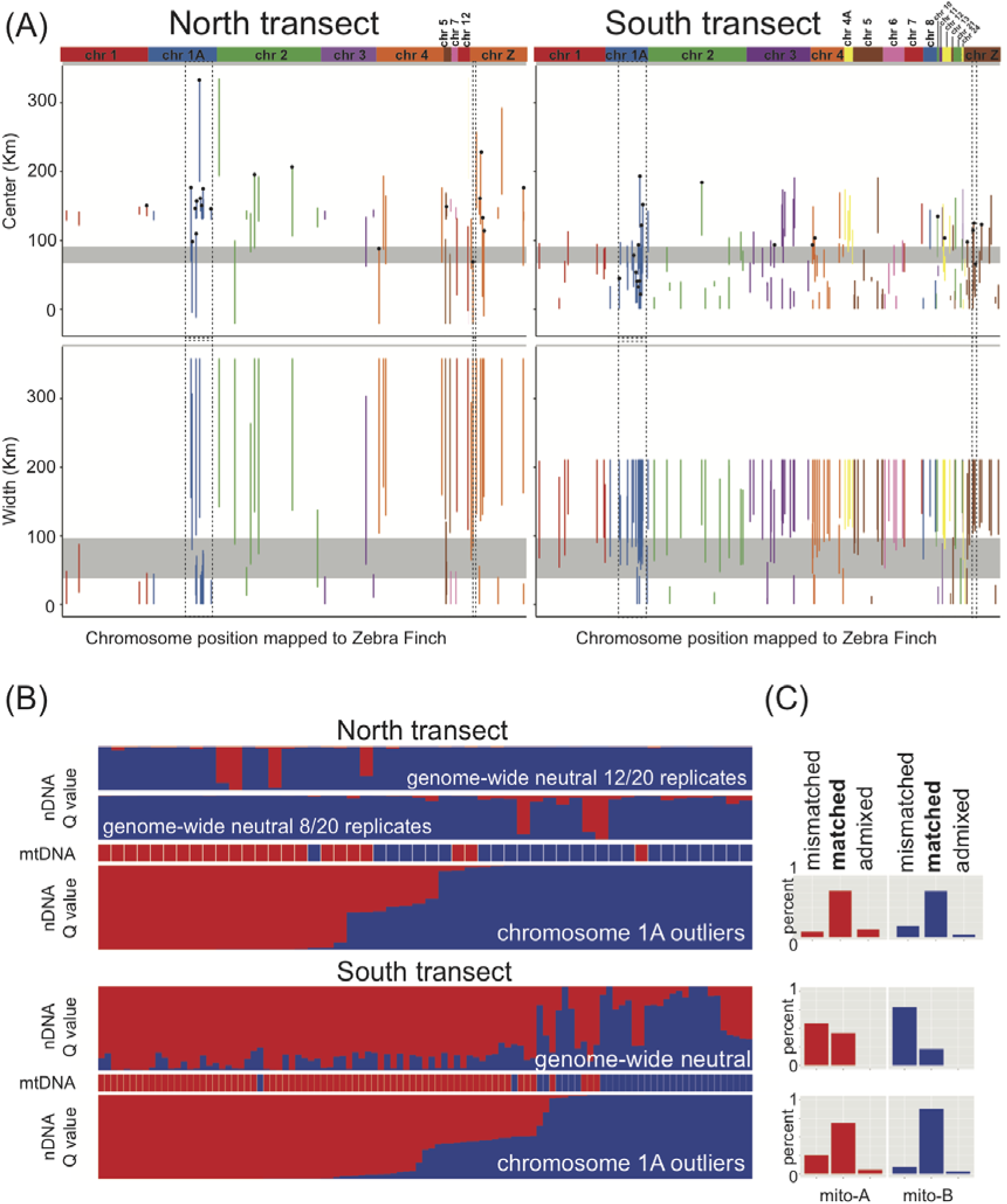
Mitolineage contact zone dynamics: (A) genome-wide levels of introgression at clinal loci, (B) individual admixture at chromosome 1A outliers and genome-wide non-outliers in north and south transect and (C) correspondence of nuclear cluster membership with mitolineage memberships in each transect for different classes of loci. (A) Confidence intervals (CIs) of centers and widths of geographic clines for northern (left panel) and southern (right panel) transects. Vertical lines represent CIs for cline parameters of nuclear clinal loci. CI lines are coloured by chromosome location. Horizontal grey bars represent CIs for cline parameters of mitochondrial clines. Loci with coloured lines that overlap the dark grey bar are inferred to have similar cline centres and/or widths to the mitochondrial clines. The position of the genomic islands of differentiation on chromosome 1A and chromosome Z are indicated by dashed rectangles. Black dots on top of coloured lines indicate outlier loci. (B) Results of STRUCTURE analyses (nDNA bar plots) showing individual probability of assignment (Q value, y-axes) to two genetic clusters (K = 2, red and blue) for 6,947 genome-wide putatively neutral loci (i.e. non-outliers located at least 100,000 bases away from any outliers) and 292 outliers located within the chromosome 1A island of differentiation. Corresponding mitolineage membership for each individual is shown with mtDNA bar plots (red-mito-A, blue-mito-B). Individuals are displayed in the order of decreasing membership in the inland cluster inferred from the outlier loci analyses. Analysis of neutral loci in the northern transect resulted in two conflicting structure patterns supported by twelve and eight replicate runs. Nuclear-mitolineage correspondence and geographic location of each individual are shown on Figs. S13-S14. (C) Percentage of individuals with nuclear variation matching their respective mitolineage (Q ≥ 0.9), with admixed nuclear ancestry (0.9 < Q > 0.1) and with nuclear-mitolineage mismatch is shown in the histograms for mito-A (red) and mito-B (blue) mitolineages. Plot for neutral loci in the north is not shown as neither of the alternative structures show corresponded to mitolineage membership.

Using STRUCTURE v.2.3.4 (Pritchard *et al.* 2000) independently for each transect, we estimated the probability of assignment of each individual (Q) to two nuclear genetic clusters (K = 2) at putatively neutral loci (i.e. 6,947 non-outliers located at least 100,000 bases away from any outliers) and chromosome 1A outliers (i.e. 292 outliers located within chromosome 1A island of differentiation). Although strong linkage of the chromosome 1A outliers violated STRUCTURE assumption of no LD among loci, similar results were obtained using assumption-free PCA (Fig. S11), and here we are using STRUCTURE Q-values to categorize individuals according to what suite of outliers they possess. Analysis of neutral loci in the northern transect resulted in two conflicting structure patterns among replicates, neither of which corresponded to mitolineage membership (Fig. 3B). The three remaining analyses assigned all individuals to two clusters, one of which prevailed in the inland, and one in the coast (Fig. 3B, Figs. S13-S14); for each analysis, assignments were consistent across all replicate runs. We classified all individuals into three groups according to the correspondence of their mitolineage with their assignment into inland or coastal cluster: matched (Q ≥ 0.9), admixed (0.9 < Q > 0.1) or mismatched (Q ≤ 0.1) (Fig. 3C). Outlier loci analyses showed that in most individuals, mitolineage matched nuclear cluster assignment (north 78%, south 75%), some individuals were admixed for both nuclear clusters (north 15%, south 20%), and a few individuals had a mitolineage that mismatched their chromosome 1A island Q-value (north 8%, south 5%). Mismatched or admixed individuals tended to be close to the contact zones between mitolineages in each transect (Suppl. Fig. S12-S14). For STRUCTURE assignments based on neutral loci in the southern transect, mitolineage matched nuclear cluster assignment in 37% of individuals, but the majority of individuals were admixed (63%) for the two nuclear clusters, and none had mitolineages mismatched with their nuclear cluster. Mismatch analysis for neutral loci in the northern transect could not be performed because its structure approximates a single homogeneous cluster without correspondence with mitolineage membership.

### Evidence for selective sweeps in the chromosome 1A genomic island of differentiation

We compared genome-wide linkage disequilibrium (LD) with that of the two chromosomes bearing significant genomic islands of differentiation. Genome-wide LD decayed quickly and reached average LD levels at a genetic distance of ~7.8 Kb between markers (Fig. 4A; Table S6). In striking contrast, chromosome 1A had the highest values of LD of all chromosomes and a substantially slower LD decay, reaching average LD at ~140 Kb (Fig. 4A; Table S6; Fig. S14). The Z chromosome also showed high levels of LD and slow decay (Fig. 4A; Table S6; Fig. S14).

**Figure 4.**
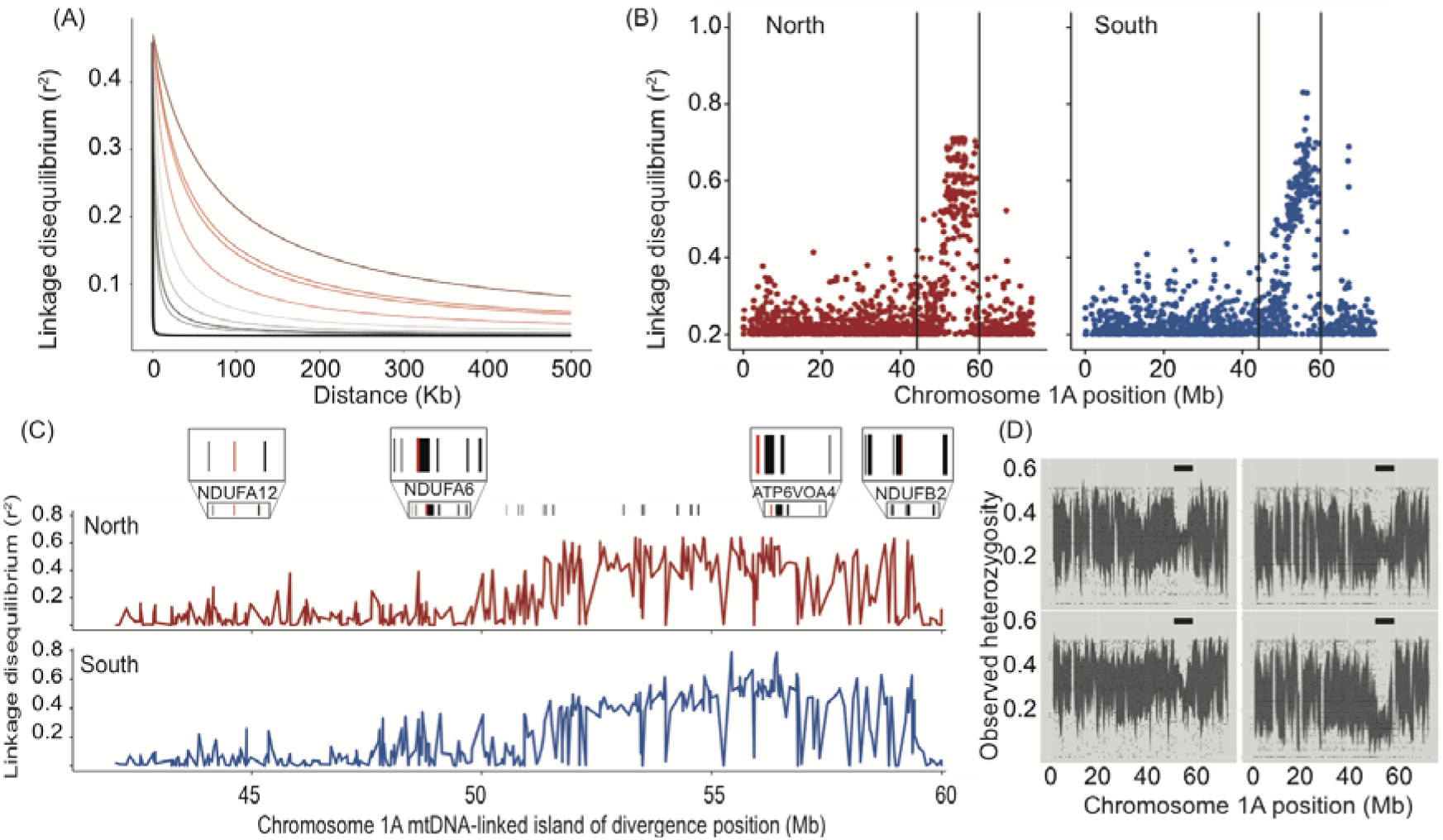
Linkage disequilibrium and genetic diversity at genomic islands of differentiation. (A) Plot of LD decay against physical distance for two chromosomes with genomic islands of differentiation, chromosome 1A (shades of red) and chromosome Z (shades of grey), and for average LD across the whole genome (thick black line on the bottom). Each line represents different mitolineage-bearing populations in each transect (from top to bottom: north mito-A-bearing, south mito-A-bearing, north mito-B-bearing and south mito-B-bearing). For similar plots of LD decay for other chromosomes see Fig. S14. (B) Mitochondrial-nuclear LD (one dot per nuclear marker) across chromosome 1A for each transect, the location of the genomic island of differentiation is indicated with vertical black lines. (C) Mitochondrial-nuclear LD within the chromosome 1A genomic island of differentiation for each transect. Lines show the transition between per nuclear marker LD values. The relative position of genes with functional annotations to mitochondrial activity are indicated with thin black bars (i.e. N-mt genes; GO term: 0005739), with protein-coding OXPHOS genes highlighted in red (GO term: 0006119). Four genome regions with high concentrations of N-mt genes including OXPHOS ones are magnified in insets. The name of the OXPHOS genes that occur within the magnified regions are indicated below the insets (see Table S5). (D) Low observed heterozygosity in chromosome 1A in different mitolineage-bearing populations in each transect. Clock-wise from top-right: north mito-A-bearing; north mito-B-bearing; south mito-A-bearing; and south mito-B-bearing. Each dot represents a locus mapped to chromosome 1A. The location of the genomic island of differentiation is indicated with horizontal black lines. To aid visualization, the grey shading shows per-SNP 95% confidence intervals within overlapping sliding windows (window size = 100 Kb, step-size = 50 Kb).

Next, we investigated the pattern of LD across chromosome 1A. Given that two neighbouring nuclear markers in strong LD with the mitochondrial genome will necessarily be in strong LD with each other, we estimated the level of mitochondrialnuclear LD in each transect (Sloan et al. 2015). Markers within the chromosome 1A genomic island of differentiation have exceptionally high levels of mitochondrial-nuclear LD compared to markers outside the island of differentiation (Fig. 4B). Linkage disequilibrium within the genomic islands of differentiation is highly heterogeneous, with regions of low mtDNA-nDNA correlations interspersed among regions of high correlations (Fig. 4C). With the present data, we cannot distinguish among the possible explanations for this heterogeneity, including occasional recombination, multiple targets of selection, ancestral polymorphism and *de novo* mutations. Different mitolineage-bearing populations in each transect showed low observed heterozygosity in the chromosome 1A genomic island of differentiation (Fig. 4D), suggesting a selective sweep (Kim & Nielsen 2004), paralleling the selective sweep previously inferred for mtDNA (Morales *et al.* 2015).

## Discussion

We used a high-density genome scan and replicated sampling to estimate genome-wide genetic differentiation and introgression between populations harbouring two deeply divergent Eastern Yellow Robin (EYR) mitolineages that occupy contrasting climates and undergo nuclear male-mediated gene flow. Inferred from comparisons with the zebra finch reference genome, genomic differentiation and restricted introgression is concentrated in a ~15.4 Mb genomic region on chromosome 1A. This genomic island of differentiation contains an overrepresentation of genes encoding protein products with mitochondrial functions (N-mt gene products). Three of these N-mt genes encode protein products of OXPHOS complex I (Fiedorczuk *et al.* 2016; Fig. S9), which might undergo long-range structural and functional interactions with three mtDNA genes that also encode components of OXPHOS complex I and were previously inferred to be under selection in EYR (Lamb *et al.*, submitted; Morales *et al.*, 2015). These results are consistent with our hypothesis that non-neutral inland-coastal mitolineage differentiation in the face of nuclear gene flow has been accompanied by non-neutral N-mt differentiation enforced by selection for mitonuclear interactions. Most loci within the detected island of differentiation on chromosome 1A have low genetic diversity and high linkage disequilibrium, as typically seen under selective sweeps. This parallels signatures of selective sweeps previously inferred in EYR mitogenomes (Morales *et al.* 2015), suggesting strong co-evolutionary dynamics. The current pattern of mitonuclear divergence in EYR is in accordance with our hypothesis of nuclear compensatory evolution, past selective sweeps and mitonuclear cointrogression of N-mt genes with mtDNA genes. We thus propose that the current pattern of geographic separation of mitolineages is maintained despite ongoing male-mediated nuclear gene flow by one or more selective mechanisms: environmental-based selection for mitonuclear co-adaption, selection against mitonuclear incompatibilities, and genomic architecture favouring mitonuclear interactions.

To our knowledge, this the first report of a genomic island of differentiation implicated in mitonuclear co-evolution. We propose that interacting N-mt and mtDNA alleles started to differentiate at the initial north-south population divergence approximately two million years ago. During this period, the fast-evolving mitogenome likely accumulated differentiation at a faster rate than the nuclear genome. Genetic signatures in mitochondrial genomes, and demographic reconstructions, suggest that large effective population sizes promoted the accumulation of mitochondrial adaptive variation that swept to fixation and was later maintained by strong purifying selection (Morales *et al.* 2015; Morales *et al.* 2017b). This may have exerted selective pressures for compensatory evolution of N-mt genes leading to selective sweeps, including in the region of the chromosome 1A island of differentiation. Under this scenario, N-mt interacting alleles introgressed together with mitogenomes (i.e. mitonuclear co-introgression), resulting in the contemporary pattern of inland-coastal divergence of N-mt and mtDNA genes. Further refinement and maintenance of mitonuclear interactions may have occurred more recently in response to ecologically-based selection across the inland-coastal climatic divide, to hybrid fitness costs from mitonuclear incompatibilities and to genomic architecture that promotes reduced recombination (see below). The scenario above is consistent with both of the general models proposed to explain the evolution of genomic island of differentiation: selective sweeps, introgression and background-selection vs. ecological-divergent selection (Cruickshank & Hahn 2014; Irwin *et al.* 2016; Nosil *et al.* 2009). Interpreting genomic islands of differentiation with genomic scan data however, remains a challenging task because many of the confounding factors that contribute to the emergence of islands cannot be controlled for with this type of data (e.g. recombination rate, mutation rate and gene density; Ravinet *et al.* 2017). Further development of EYR genomic resources is needed to test relative vs. absolute metrics of genetic differentiation and better distinguish alternative models of island formation (Cruickshank & Hahn 2014). Likewise, exploring patterns of chromosome 1A genetic diversity and differentiation across a range of passerines species is needed to understand the relative contribution of different mechanisms (e.g. recombination and mutation rates) to island formation over evolutionary timescale (Burri *et al.* 2015; Irwin *et al.* 2016).

Our data suggest that natural selection plays a major role in EYR’s mitonuclear divergence in two main ways: selection against mitonuclear genetic incompatibilities (intrinsic barriers) and selection for locally adapted mitonuclear interactions (extrinsic barriers) (Bierne *et al.* 2011; Burton & Barreto 2012; Burton *et al.* 2013; Hill 2015; Qvarnström *et al.* 2016). Nuclear admixture and mitonuclear mismatch at chromosome 1A outlier loci are rarer than in genome-wide neutral loci, and admixed or mismatched individuals are predominantly found within the contact zones between mitolineages. This suggest that selection acts against the formation of hybrids with incompatible mitonuclear combinations via the formation of intrinsic reproductive barriers (e.g. assortative mating). On the other hand, strong fine-spatial-scale (20 - 40 km) separation of mitolineages across the climatic divide, despite birds’ capability to travel several km per day (Debus & Ford 2012), suggests that extrinsic reproductive barriers prevents settling of dispersing birds in the environment inappropriate for their mitolineage. Likewise, strong correlations of allelic frequencies of outlier loci across the environmental gradient between northern and southern nuclear backgrounds supports a scenario of replicated environmentally-based divergence of N-mt genes.

Both of the above mechanisms for maintaining mitonuclear divergence may be supported by a favourable genomic architecture. High LD in the chromosome 1A genomic region enriched for N-mt genes suggests that reduced rates of recombination could prevent formation of mitonuclear incompatibilities by promoting the co-inheritance of matching combinations of interacting N-mt and mtDNA genes. Accordingly, low recombination could allow locally co-adapted alleles to sustain nuclear-nuclear and mitochondrial-nuclear interactions (Lindtke & Buerkle 2015; Ortiz-Barrientos *et al.* 2016; Sunnucks *et al.* 2017; Yeaman 2013). The inferred location of the chromosome 1A island of differentiation overlaps the position of the chromosome’s centromere in zebra finch (Warren *et al.* 2010), and centromeres are known to undergo reduced recombination and contribute disproportionately to genetic divergence (Butlin 2005; Kawakami *et al.* 2014). Intriguingly, clustered patterns of genetic differentiation have been identified at similar genomic locations on chromosome 1A in at least two other passerine birds that also harbour deep mitochondrial divergences: greenish warblers, *Phylloscopus trochiloides* (Irwin *et al.* 2016), and flycatchers, *Ficedula albicollis and F. hypoleuca* (Ellegren *et al.* 2012). This suggests that genomic architecture of these regions may enforce cold-spots of recombination over large evolutionary timescales, in agreement with the observation that the landscape of recombination in birds is highly conserved (Singhal *et al.* 2015). Also, these findings suggest that this genomic region may contain an ancient configuration of N-mt genes commonly involved in mitonuclear divergence. Given that some of these N-mt gene products occur in functionally-related regions of complex I (Fig. S9), structural or mechanistic complementarity between them may have driven the mitonuclear co-evolution over long evolutionary time-scales, but sequencing data for these N-mt genes are need to further explore potential functional relationships between their products.

We identified a second genomic island of differentiation in the Z sex chromosome. The Z chromosome, compared to all autosomes, had significantly higher levels of genetic differentiation between different mitolineage-bearing populations in each transect, except for chromosomes 1A and 23 (pairwise Wilcoxon test, P-value for all significant tests ≤ 0.007). This is consistent with the widespread observation that sex chromosomes often show elevated divergence between differentiated lineages (Mank *et al.* 2010; Qvarnström & Bailey 2009). Moreover, stronger genetic incompatibilities in hybrid heterogametic (ZW) females than homogametic males (ZZ) (i.e. Haldane’s Rule; Haldane 1922) could lead to reduced female-mediated Z-linked and mitochondrial gene flow compared to male-mediated nuclear gene flow (Beekman *et al.* 2014). Therefore, Haldane’s Rule could explain how EYR experiences inland-coastal male-mediated gene flow, despite females being the most dispersing sex (Debus & Ford 2012; Harrisson *et al.* 2012). Alternatively, the Z chromosome could contain barrier loci directly related to species recognition and mitonuclear metabolism (Hill & Johnson 2013; Qvarnström & Bailey 2009). A previous study found limited evidence for sexual dimorphism in EYR colouration, but identified small but significant colour differences between inland and coastal individuals (Morales *et al.* 2017a). Haldane’s Rule and mitonuclear sexual selection predict different patterns of genetic differentiation along the Z chromosome. Under Haldane’s Rule, genetically differentiated loci should be spread through the Z chromosome, while under mitonuclear sexual selection, differentiation should be concentrated in genomic regions containing N-mt genes. The present data on EYR seem to show both patterns: widespread distribution of outliers along the Z chromosome, but also a narrow island of differentiation. However, it is important to note that our loci mapped to Z-chromosome in the zebra finch mapped to different regions in the collared flycatcher, *F. albicollis*, genome (Ellegren *et al.* 2012) (see Methods), reducing our ability to interpret the pattern of differentiation (Fig. S15).

In summary, our study contributes to the mounting evidence that mitonuclear co-evolution is an important mechanism of population divergence (Bar-Yaacov *et al.* 2015; Baris *et al.* 2017; Boratyński *et al.* 2016; Burton & Barreto 2012; Burton *et al.* 2013; Ellison & Burton 2006; Gagnaire *et al.* 2012; Hill 2015; Hill 2016; Sambatti *et al.* 2008). Future studies should focus on the genomic and phenotypic basis of EYR mitonuclear co-evolution by expanding EYR genomic resources (e.g. whole genome and transcriptome sequencing) and collecting data on fitness proxies related to mitonuclear functioning. These efforts will enable us to answer important questions about genomic landscape of differentiation (Cruickshank & Hahn 2014; Irwin *et al.* 2016; Nosil *et al.* 2009), including the contribution of linked selection and heterogeneous levels of gene flow and recombination (Roux *et al.* 2016; Schrider *et al.* 2016), the age of beneficial allele combinations (Smith *et al.* 2016), the genomic mechanisms to maintain mitonuclear interactions despite gene flow (Schwander *et al.* 2014; Sunnucks *et al.* 2017), the relationships between mitochondrial metabolism and environment (Boratyński *et al.* 2016; McFarlane *et al.* 2016; Stier *et al.* 2014; Toews *et al.* 2014), and fitness consequences of mitonuclear genetic incompatibilities and mitonuclear co-adaptation (Burton & Barreto 2012; Burton *et al.* 2013; Hill 2015; Hill 2016).

## Methods

### Samples and mitolineage identification

We determined the EYR mitolineage for 407 individuals (N_mito-A_ = 262; N_mito-B_ = 145) using ND2 sequences (Table S1). DNeasy Blood and Tissue Kit (Qiagen, Germany) was used to extract DNA. PCRs were performed following Pavlova *et al.* (2013) and sequenced commercially (Macrogen, Korea).

### Climatic variables

Values for two BIOCLIM variables, maximum temperature of the warmest month (BIOCLIM 5) and minimum precipitation of the driest month (BIOCLIM 14), were extracted for each sampling location and ~350 random points (equally spaced at 0.02 degrees in each transect) using the R package *raster* (Hijmans 2014; Hijmans *et al.* 2005; R Development Core Team 2014). Random points were used to represent background climatic variation in each transect by fitting a linear model of climatic variation against a geographic variable (longitude in the north transect and latitude in the south transect) with the function *lm* in R. We quantified the Pearson’s r correlation between mitolineage membership and climatic variation using ND2 samples with the function *cor* in R.

### Sequencing, genotyping and mapping

We genotyped samples using the reduced-representation approach implemented in DarTseq (Diversity Arrays Technology, Australia) (see supplementary methods for details). Samples were digested using a combination of *Pst*I and *Nsp*I enzymes following Kilian *et al.* (2012). Reduced-representation libraries were prepared for Illumina sequencing including a varying length barcode region (including 63 technical replicates). Libraries were sequenced with 77 single-read sequencing cycles on Illumina Hiseq2000. Sequencing reads were processed using proprietary DArT P/L’s analytical pipelines (see Supplementary file for details).

A total of 68,258 DArT-tags (60 bp each after trimming barcodes) containing 97,070 SNP markers were obtained (raw data are deposited to Figshare doi: 10.6084/m9.figshare.5072143). We retained 60,444 SNPs that had a call rate of at least 80%. We did not filter SNPs by Minor Allele Frequency (MAF) in the main dataset after confirming that results do not change after filtering-out SNPs with MAF < 5% (data not shown).

We approximated the genomic position of each SNP by mapping DArT-tags to the reference genome of the zebra finch *Taeniopygia guttata,* taeGut3.2.4 (Warren *et al.* 2010) using BLASTn v.2.3.0 (Altschul *et al.* 1990; Camacho *et al.* 2009). Only unique high-quality hits were considered. Considering only unique hits considerably increases our confidence in the mapping. However, we were able to map only 55% of the DArT-tags with this method, a limitation of not having a reference genome for the EYR. Nonetheless, markers were distributed over all autosomes and the Z chromosome except the micro-chromosomes 16, LG2 and LG5 and there was a strong positive correlation between the number of mapped tags and chromosome size, indicating that tags mapped uniformly across the zebra finch genome (Pearson’s r = 0.99; Fig. S16). We were unable to assess the coverage of the female-specific W-chromosome because it is absent from the zebra finch reference genome. We were unstable to place a few hundred outlier loci in the genome (Fig. S17).

We explored potential conflicts in our mapping by comparing results from the zebra finch reference genome (Warren *et al.* 2010) to the collared flycatcher reference genome (Ellegren *et al.* 2012). Results show that DArT-tags mapped to very similar genomic positions in both genomes for most of the chromosomes, including the region of the chromosome 1A genomic island of differentiation (Fig. S15). The position of DArT-tags that mapped to the sex chromosome Z however, was substantially different in the two reference genomes, highlighting potential conflicts with our mapping in this chromosome. The zebra finch genome is our preferred reference given its more advanced level of annotation.

### Genetic differentiation between different mitolineage-bearing populations in each transect and identification of outliers

We summarised genetic structure of all samples with Principal Component Analysis (PCA) using the *dudi.pca* function in Adegenet 2.0.1 (Jombart 2008). We estimated genetic differentiation between different mitolineage-bearing populations for each transect and identified outlier loci (i.e. loci of extreme differentiation compared to the average) with three different methods:

#### 1) *F*_ST_ *-outlier detection at fine spatial scales*

We measured per-SNP genetic differentiation (Weir and Cockerham’s *F*_ST_) between different mitolineage-bearing populations in each transect with the *diffCalc* function of the R package *DiveRsity* and considered the upper 1% values as outlier loci (Keenan *et al.* 2013; Weir & Cockerham 1984). We limited the samples to individuals within a 40-km radius from the mid-point of the contact zone (11 north-mito-A, 20 north-mito-B; 20 south-mito-A and 20 south-mito-B). We assumed that within an 80-km-wide region the effect of genetic drift is negligible, given that EYR disperses 2-25 km per generation (Debus & Ford 2012; N. Amos, unpublished analysis of Australian Bird and Bat Banding Scheme data).

#### 2) Correlations between nuclear loci and mitolineage membership with BayScEnv

We used BayeScEnv, which detects loci that depart from neutral expectations (outliers) based on their *F*_ST_ values and associations of allele frequencies with environmental variables, while correcting for the confounding effects of population history. SNPs are assigned either to a neutral model or one of two non-neutral models: the environmental correlation model and the locus-specific model (Villemereuil & Gaggiotti 2015). Samples were grouped into populations according to their geographic location (Fig. S18). We used the mitolineage membership of each population as a binomial environmental variable (mito-A coded 1 and mito-B coded -1). We used twenty pilot runs of 4,000 iterations with a burn-in of 80,000 iterations and samples taken every 10 steps and a False Discovery Rate (FDR) significance threshold of 5%. Convergence of every run was confirmed using the R package *coda* (Plummer *et al.* 2006). We assigned equal prior probability to both non-neutral models, but only the model based on mitolineage membership produced outliers.

#### 3) Principal component analysis

We estimated genetic differentiation across each transect without assuming any kind of prior grouping, using the program PCAdapt (Duforet-Frebourg *et al.* 2016). PCAdapt uses a hierarchical Bayesian model to determine population structure with latent factors (K, analogue to PCA axes) and identify outlier loci that contribute disproportionately to explaining each of the K factors. Among the important differences from other methods are that PCAdapt does not rely on *F*_ST_ estimates, does not require classification of individuals into populations, performs well across a range of demographic models, and it is agnostic to environmental information (or mitolineage membership in our case). An initial inspection of 76 K factors revealed that while the first seven K’s explain most of the genetic variation, only K1 differentiated well between different mitolineage-bearing populations in both transects (Fig. S19). Accordingly, we performed the analysis with K=2 and extracted outliers along K1.

### Allelic frequency correlations

To test whether alleles from outlier loci were segregating in the same direction in both divergent nuclear genomic backgrounds, we performed a correlation analysis of allelic frequencies between different mitolineage-bearing populations in the northern and southern transects: north-mito-A versus south-mito-A and north-mito-B versus south-mito-B. We compared outlier loci correlations with correlations drawn from randomly selected loci across the rest of the genome (i.e. random distribution). We included outlier loci that were identified in both transects by at least one of the methods (N = 249). The random distribution was calculated with 240 random sets of non-outlier loci, each containing 249 loci. We tested if allelic correlations of outlier loci were greater than expected at random with a t-test in R.

### Identification of genomic islands of differentiation

Hidden Markov Models (HMM) are useful to identify genomic regions that contain contiguous SNPs of high differentiation without having to rely on methods that require defining arbitrary sliding windows sizes (Hofer *et al.* 2012). HMM assumes that genetic differentiation changes across the genome between hidden states and assigns each SNP to a given state level and identifies state transitions. We defined three hidden states of genetic differentiation - low, intermediate and high - and considered genomic islands of differentiation as clusters of contiguous SNPs of high differentiation. We used the cumulative distribution function of q-values from the BayeScEnv analysis with mitolineage membership as input. We corrected for multiple-testing correction with a FDR significance threshold of 1% and used a reduced datset of 27,912 SNPs with a call rate higher than 90% and a MAF > 10% to avoid rare variants following Marques *et al.* (2016). To overcome low marker density for some chromosomes, we modelled hidden state changes across the entire genome. This decision did not bias our results because we did not identify any significant state transition between chromosomes. The analysis was performed with the R package *HiddenMarkov* (Harte 2015).

### Functional significance of candidates for mitochondrial-nuclear interactions

We counted zebra finch genes with functional annotations for mitochondrial activity (i.e. N-mt genes; GO term: 0005739) extracted from ENSEMBL (accessed November 2015, Cunningham *et al.* 2015), and a subset of those encoding supernumerary subunits and assembly factors for OXPHOS complexes (i.e. OXPHOS genes; GO term: 0006119) extracted from the Kyoto Encyclopaedia of Genes and Genomes (KEGG, accessed June 2015; Kanehisa & Goto 2000). We performed the counts within each of the genomic islands of differentiation and equal-sized regions across the whole genome, avoiding islands (i.e. random distribution). For each island of differentiation, we computed the probability that the observed count was significantly higher than the random distribution with the *t.test* function in R.

We mapped the protein products of three nuclear OXPHOS genes located within the chromosome 1A genomic island of differentiation and of three EYR mitochondrial OXPHOS genes previously suggested to be under divergent selection (Lamb *et al.* submitted; Morales *et al.* 2015) onto the 4.2 Å resolution published structure of the homologous cryo-EM bovine mitochondrial complex I (Zhu *et al.* 2016) in the molecular graphics program UCSF Chimera (Pettersen *et al.* 2004).

### Geographic cline analysis

We examined how allelic frequencies change as a function of geographic distance across each transect. We used the R package *hzar* (Derryberry *et al.* 2014), which implements a MCMC approach to fit allelic frequency data to cline models (Barton & Gale 1993). We used individual nuclear allele frequencies (0, 0.5 or 1) and mitochondrial haplotypes (mito-A coded 0 and mito-B coded 1) to fit the cline models. We reduced linkage between markers and computation times by randomly selecting one SNP every 100 Kb along each chromosome (2,494 nuclear loci + mtDNA = 2,495 data points per transect). For each transect, we projected the geographic location of each individual on a unidimensional axis (Fig. S10). Then, we calculated the distance of each projected location to a common geographic point. For the southern transect we removed the samples furthest from the contact zone (lower left corner in the dashed south square in Fig. 1A) because the geographic gap between these samples and the rest of the samples was very large and could bias the cline-fitting, resulting in a shorter unidimensional axis (Fig. S10). We allowed a buffer distance of +/- 30 km at both ends of each transect following Derryberry *et al.* (2014) to ensure that all samples within each transect were fitted in the cline models. We fitted three independent models, all of which estimate cline centre (i.e. geographic location of the maximum allelic frequency change) and width (defined as 1/maximum slope; Derryberry *et al.* 2014) but differed in the way they fitted cline tails (i.e. the exponential decay curve). Model I fitted fixed cline parameters and no tails. Model II fitted variable cline parameters and no tails. Model III fitted variable cline parameters and two mirroring tails. Models were compared against a neutral cline model (flat, no clinal variation) with the Akaike information criterion (AICc), and this was also used to choose the best clinal model for each SNP.

### Admixture analyses

We estimated the probability of assignment (Q) of each individual to two nuclear genetic clusters using the admixture model with correlated allele frequencies implemented in STRUCTURE v.2.3.4 (Pritchard *et al.* 2000) using two different datasets. The first dataset included 6,947 genome-wide neutral loci: we first removed all outlier loci and all loci within 100,000 bases of any outliers, and then we randomly sampled one nonoutlier locus every 100,000 bases along the entire genome. The second dataset included 292 outlier loci located within the chromosome 1A island of differentiation. Separate analyses were run for northern (50 individuals) and southern (103 individuals) transects. To account for unequal representation of individuals in each of the source populations, population-specific ancestry priors were used (Wang 2017). For each marker set we performed one STRUCTURE analysis per transect assuming Kmax=2. Twenty replicates of 200,000 burn-in iterations followed by 1,000,000 iterations run for each K. Runs were summarized using web server CLUMPAK (Kopelman *et al.* 2015).

### Genetic diversity and Linkage Disequilibrium (LD)

We calculated observed heterozygosity as a proxy for per-locus genetic diversity using the *basicStats* function of the R package *DiveRsity* (Keenan *et al.* 2013). We calculated pairwise LD per chromosome and the rate of LD decay as a function of physical distance across the entire genome (all chromosomes together) and independently for two chromosomes with genomic islands of differentiation, 1A and Z. For pairs of different mitolineage-bearing populations in each transect we first estimated LD (r^2^) between each pair of SNP markers from the reduced SNP dataset within each chromosome with PLINK (Purcell *et al.* 2007). We then calculated the overall decay of LD against distance following Hilland Weir (1988) as implemented in Marroni *et al.* (2011).

We estimated mitochondrial-nuclear LD with a custom perl script (electronic supplementary material, file S1 from Sloan *et al.* 2015). This method calculates the per-locus correlation between nuclear and mitochondrial alleles by testing one randomly-selected nuclear allele against its mitolineage membership in each transect, and assigns statistical significance with a Fisher’s exact test. P-values were calculated by Monte Carlo simulations (1 x 10^6^ replicates) and adjusted with a FDR significance threshold of 5%. We used a reduced dataset of 27,912 SNPs with a call rate higher than 90% and a MAF > 10% to avoid rare variants following Sloan *et al.* (2015)

### Data availability

All datasets were deposited in Figshare doi: 10.6084/m9.figshare.5072143 (temporary link until publication: https://figshare.com/s/3476e05a55632554d37a) and comprised (1) a table with information for every individual including geo-climatic data, ND2 sequences and DArT genotypes; (2) unfiltered SNP dataset with 97,070 SNP markers; and (3) BLAST results for reference genomes.

R custom scripts were deposited in GitHub: https://github.com/hmoral/EYR_DArT and comprised (1) script for HMM analyses; (2) script for allelic frequency correlations; (3) script for geographic cline analyses; and (4) script for N-mt gene enrichment test analyses.

## Acknowledgments

HM was supported by the Holsworth Wildlife Research Endowment (2012001942) and Stuart Leslie Bird Research Award from BirdLife Australia, PhD scholarships from Monash University and the Department of Public Education (SEP) of the Mexican Government, and a Monash Postgraduate Publication Award. Other funding came from Monash internal sources. Genomic analyses were undertaken at the Monash high-performance computing facility. Field samples were collected under scientific research permits issued by the Victorian Department of Environment and Primary Industries (numbers 10007165, 10005919 and 10005514), New South Wales Office of Environment and Heritage (SL100886), using bands issued by the Australian Bird and Bat Banding Scheme. We are grateful to Leo Joseph, Robert Palmer, Holly Sitters and Christine Connelly for providing genetic samples. Anders Gonçalves da Silva, David Marques and Victor Soria-Carrasco provided valuable inputs regarding data analysis, Leo Joseph on EYR evolution, and Jonci Wolf on functional properties of the mitonuclear candidates. We thank Scott Edwards, Mike Webster, Lynna Kvistad and Stephanie Falk for comments on earlier versions of the manuscript

## References

Allen JF (2003) The function of genomes in bioenergetic organelles. Philosophical Transactions of the Royal Society of London B: Biological Sciences 358, 19–38.

Altschul SF, Gish W, Miller W, Myers EW, Lipman DJ (1990) Basic local alignment search tool. Journal of molecular biology 215, 403–410.

Angerer H, Radermacher M, Mankowska M, et al. (2014) The LYR protein subunit NB4M/NDUFA6 of mitochondrial complex I anchors an acyl carrier protein and is essential for catalytic activity. Proc Natl Acad Sci U S A 111, 5207–5212.

Arnqvist G, Dowling DK, Eady P, et al. (2010) Genetic architecture of metabolic rate: environment specific epistasis between mitochondrial and nuclear genes in an insect. Evolution 64, 3354–3363.

Ballard JWO, Pichaud N (2014) Mitochondrial DNA: more than an evolutionary bystander. Functional Ecology 28, 218–231.

Bar-Yaacov D, Blumberg A, Mishmar D (2012) Mitochondrial-nuclear co-evolution and its effects on OXPHOS activity and regulation. Biochimica et Biophysica Acta (BBA)-Gene Regulatory Mechanisms 1819, 1107–1111.

Bar-Yaacov D, Hadjivasiliou Z, Levin L, et al. (2015) Mitochondrial involvement in vertebrate speciation? The case of mito-nuclear genetic divergence in chameleons. Genome biology and evolution 7, 3322–3336.

Baris TZ, Wagner DN, Dayan DI, et al. (2017) Evolved genetic and phenotypic differences due to mitochondrial-nuclear interactions. PLoS genetics 13, e1006517.

Barton NH, Gale KS (1993) Genetic analysis of hybrid zones. In: Hybrid zones and the evolutionary process (ed. Harrison R), pp. 13–45. Oxford University Press, New York.

Beck EA, Thompson AC, Sharbrough J, Brud E, Llopart A (2015) Gene flow between *Drosophila yakuba* and *Drosophila santomea* in subunit V of cytochrome c oxidase: A potential case of cytonuclear cointrogression. Evolution 69, 1973–1986.

Beekman M, Dowling DK, Aanen DK (2014) The costs of being male: are there sex-specific effects of uniparental mitochondrial inheritance? Philosophical Transactions of the Royal Society B: Biological Sciences 369, 20130440.

Bierne N, Welch J, Loire E, Bonhomme F, David P (2011) The coupling hypothesis: why genome scans may fail to map local adaptation genes. Molecular Ecology 20, 2044–2072.

Boratyński Z, Ketola T, Koskela E, Mappes T (2016) The Sex Specific Genetic Variation of Energetics in Bank Voles, Consequences of Introgression? Evolutionary Biology 43, 37–47.

Burri R, Nater A, Kawakami T, et al. (2015) Linked selection and recombination rate variation drive the evolution of the genomic landscape of differentiation across the speciation continuum of *Ficedula flycatchers*. Genome Research 25, 1656–1665.

Burton RS, Barreto FS (2012) A disproportionate role for mtDNA in Dobzhansky–Muller incompatibilities? Molecular Ecology 21, 4942–4957.

Burton RS, Pereira RJ, Barreto FS (2013) Cytonuclear Genomic Interactions and Hybrid Breakdown. Annual Review of Ecology, Evolution, and Systematics 44, 281–302.

Butlin RK (2005) Recombination and speciation. Molecular Ecology 14, 2621–2635.

Camacho C, Coulouris G, Avagyan V, et al. (2009) BLAST+: architecture and applications. BMC Bioinformatics 10, 1.

Cruickshank TE, Hahn MW (2014) Reanalysis suggests that genomic islands of speciation are due to reduced diversity, not reduced gene flow. Molecular Ecology 23, 3133–3157.

Cunningham F, Amode MR, Barrell D, et al. (2015) Ensembl 2015. Nucleic acids research 43, D662–D669.

Das J (2006) The role of mitochondrial respiration in physiological and evolutionary adaptation. BioEssays 28, 890–901.

Debus S, Ford H (2012) Responses of Eastern Yellow Robins *Eopsaltria australis* to translocation into vegetation remnants in a fragmented landscape. Pacific Conservation Biology 18, 194–202.

Derryberry EP, Derryberry GE, Maley JM, Brumfield RT (2014) HZAR: hybrid zone analysis using an R software package. Molecular ecology resources 14, 652–663.

Dowling DK, Friberg U, Lindell J (2008) Evolutionary implications of non-neutral mitochondrial genetic variation. Trends in Ecology and Evolution 23, 546–554.

Duforet-Frebourg N, Luu K, Laval G, Bazin E, Blum MG (2016) Detecting genomic signatures of natural selection with principal component analysis: application to the 1000 Genomes data. Molecular Biology and Evolution 33, 1082–1093.

Ellegren H, Smeds L, Burri R, et al. (2012) The genomic landscape of species divergence in Ficedula flycatchers. Nature 491, 756–760.

Ellison CK, Burton RS (2006) Disruption of mitochondrial function in interpopulation hybrids of *Tigriopus californicus*. Evolution 60, 1382–1391.

Fiedorczuk K, Letts JA, Degliesposti G, et al. (2016) Atomic structure of the entire mammalian mitochondrial complex I. Nature 538, 406–410.

Gagnaire P-A, Normandeau E, Bernatchez L (2012) Comparative genomics reveals adaptive protein evolution and a possible cytonuclear incompatibility between European and American Eels. Molecular Biology and Evolution 29, 2909–2919.

Haldane JB (1922) Sex ratio and unisexual sterility in hybrid animals. Journal of genetics 12, 101–109.

Harrison RG, Larson EL (2016) Heterogeneous genome divergence, differential introgression, and the origin and structure of hybrid zones. Molecular Ecology.

Harrisson KA, Pavlova A, Amos JN, et al. (2012) Fine-scale effects of habitat loss and fragmentation despite large-scale gene flow for some regionally declining woodland bird species. Landscape ecology 27, 813–827.

Harte D (2015) HiddenMarkov: Hidden Markov Models. R package version 1.8-4. Statistics Research Associates, Wellington. <u>http://homepages.maxnet.co.nz/davidharte/SSLib/</u>

Hijmans RJ (2014) raster: Geographic data analysis and modeling R package version 2.3-12.

Hijmans RJ, Cameron SE, Parra JL, Jones PG, Jarvis A (2005) Very high resolution interpolated climate surfaces for global land areas. International journal of climatology 25, 1965–1978.

Hill GE (2015) Mitonuclear ecology. Molecular Biology and Evolution 32, 1917–1927.

Hill GE (2016) Mitonuclear coevolution as the genesis of speciation and the mitochondrial DNA barcode gap. Ecology and evolution 22, 5831–5842.

Hill GE, Johnson JD (2013) The mitonuclear compatibility hypothesis of sexual selection. Proceedings of the Royal Society of London B: Biological Sciences 280, 20131314.

Hill W, Weir B (1988) Variances and covariances of squared linkage disequilibria in finite populations. Theoretical population biology 33, 54–78.

Hoban S, Kelley JL, Lotterhos KE, et al. (2016) Finding the Genomic Basis of Local Adaptation: Pitfalls, Practical Solutions, and Future Directions. The American naturalist 188, 379–397.

Hoekstra LA, Siddiq MA, Montooth KL (2013) Pleiotropic Effects of a Mitochondrial–Nuclear Incompatibility Depend upon the Accelerating Effect of Temperature in *Drosophila*. Genetics 195, 1129–1139.

Hofer T, Foll M, Excoffier L (2012) Evolutionary forces shaping genomic islands of population differentiation in humans. BMC genomics 13, 1.

Horan MP, Gemmell NJ, Wolff JN (2013) From evolutionary bystander to master manipulator: the emerging roles for the mitochondrial genome as a modulator of nuclear gene expression. European Journal of Human Genetics 21, 1335–1337.

Irwin DE, Alcaide M, Delmore KE, Irwin JH, Owens GL (2016) Recurrent selection explains parallel evolution of genomic regions of high relative but low absolute differentiation in a ring species. Molecular Ecology 25, 4488–4507.

James JE, Piganeau G, Eyre Walker A (2016) The rate of adaptive evolution in animal mitochondria. Molecular Ecology 25, 67–78.

Jombart T (2008) adegenet: a R package for the multivariate analysis of genetic markers. Bioinformatics 24, 1403–1405.

Jones FC, Grabherr MG, Chan YF, et al. (2012) The genomic basis of adaptive evolution in threespine sticklebacks. Nature 484, 55–61.

Kanehisa M, Goto S (2000) KEGG: kyoto encyclopedia of genes and genomes. Nucleic acids research 28, 27–30.

Kawakami T, Smeds L, Backström N, et al. (2014) A high density linkage map enables a second generation collared flycatcher genome assembly and reveals the patterns of avian recombination rate variation and chromosomal evolution. Molecular Ecology 23, 4035–4058.

Keenan K, McGinnity P, Cross TF, Crozier WW, Prodöhl PA (2013) diveRsity: An R package for the estimation and exploration of population genetics parameters and their associated errors. Methods in Ecology and Evolution 4, 782–788.

Kilian A, Wenzl P, Huttner E, et al. (2012) Diversity arrays technology: a generic genome profiling technology on open platforms. Data Production and Analysis in Population Genomics: Methods and Protocols, 67–89.

Kim Y, Nielsen R (2004) Linkage disequilibrium as a signature of selective sweeps. Genetics 167, 1513–1524.

Kopelman NM, Mayzel J, Jakobsson M, Rosenberg NA, Mayrose I (2015) Clumpak: a program for identifying clustering modes and packaging population structure inferences across K. Molecular ecology resources 15, 1179–1191.

Lamb A, Gan H, Greening C, et al. (submitted) Climate-driven mitochondrial selection: a test in Australian birds. Molecular Ecology.

Lindtke D, Buerkle CA (2015) The genetic architecture of hybrid incompatibilities and their effect on barriers to introgression in secondary contact. Evolution 69, 1987–2004.

Lowell BB, Spiegelman BM (2000) Towards a molecular understanding of adaptive thermogenesis. Nature 404, 652–660.

Mank JE, Nam K, Ellegren H (2010) Faster-Z evolution is predominantly due to genetic drift. Molecular Biology and Evolution 27, 661–670.

Marques DA, Lucek K, Meier JI, et al. (2016) Genomics of rapid incipient speciation in sympatric threespine stickleback. PLoS Genet 12, e1005887.

Marroni F, Pinosio S, Zaina G, et al. (2011) Nucleotide diversity and linkage disequilibrium in *Populus nigra* cinnamyl alcohol dehydrogenase (CAD4) gene. Tree genetics & genomes 7, 1011–1023.

McFarlane SE, Sirkiä PM, Ålund M, Qvarnström A (2016) Hybrid Dysfunction Expressed as Elevated Metabolic Rate in Male Ficedula Flycatchers. PLoS One 11, e0161547.

Meiklejohn CD, Holmbeck MA, Siddiq MA, et al. (2013) An incompatibility between a mitochondrial tRNA and its nuclear-encoded tRNA synthetase compromises development and fitness in *Drosophila*. PLoS genetics 9, e1003238.

Morales HE, Pavlova A, Joseph L, Sunnucks P (2015) Positive and purifying selection in mitochondrial genomes of a bird with mitonuclear discordance. Molecular Ecology 24, 2820–2837.

Morales HE, Pavlova A, Sunnucks P, et al. (2017a) Neutral and selective drivers of colour evolution in a widespread Australian passerine. Journal of Biogeography 44, 522–536.

Morales HE, Sunnucks P, Joseph L, Pavlova A (2017b) Perpendicular axes of differentiation generated by mitochondrial introgression. Molecular Ecology.

Nachman MW, Payseur BA (2012) Recombination rate variation and speciation: theoretical predictions and empirical results from rabbits and mice. Philosophical Transactions of the Royal Society B: Biological Sciences 367, 409–421.

Noor MA, Bennett SM (2009) Islands of speciation or mirages in the desert? Examining the role of restricted recombination in maintaining species. Heredity 103, 439–444.

Nosil P, Funk DJ, Ortiz-Barrientos D (2009) Divergent selection and heterogeneous genomic divergence. Molecular Ecology 18, 375–402.

Ortiz-Barrientos D, Engelstädter J, Rieseberg LH (2016) Recombination Rate Evolution and the Origin of Species. Trends in Ecology & Evolution 31, 226–236.

Osada N, Akashi H (2012) Mitochondrial-nuclear interactions and accelerated compensatory evolution: evidence from the primate cytochrome C oxidase complex. Molecular Biology and Evolution 29, 337–346.

Ostergaard E, Rodenburg RJ, van den Brand M, et al. (2011) Respiratory chain complex I deficiency due to NDUFA12 mutations as a new cause of Leigh syndrome. J Med Genet 48, 737–740.

Pavlova A, Amos JN, Joseph L, et al. (2013) Perched at the mito nuclear crossroads: divergent mitochondrial lineages correlate with environment in the face of ongoing nuclear gene flow in an australian bird. Evolution 67, 3412–3428.

Payseur BA, Rieseberg LH (2016) A Genomic Perspective on Hybridization and Speciation. Molecular Ecology.

Pettersen EF, Goddard TD, Huang CC, et al. (2004) UCSF Chimera—a visualization system for exploratory research and analysis. Journal of computational chemistry 25, 1605–1612.

Plummer M, Best N, Cowles K, Vines K (2006) CODA: Convergence diagnosis and output analysis for MCMC. R news 6, 7–11.

Pritchard JK, Stephens M, Donnelly P (2000) Inference of population structure using multilocus genotype data. Genetics 155, 945–959.

Purcell S, Neale B, Todd-Brown K, et al. (2007) PLINK: a tool set for whole-genome association and population-based linkage analyses. American Journal of Human Genetics 81, 559–575.

Qvarnström A, Ålund M, McFarlane SE, Sirkiä PM (2016) Climate adaptation and speciation: particular focus on reproductive barriers in *Ficedula flycatchers*. Evolutionary Applications 9, 119–134.

Qvarnström A, Bailey RI (2009) Speciation through evolution of sex-linked genes. Heredity 102, 4–15.

R Development Core Team (2014) R: A language and environment for statistical computing. R Foundation for Statistical Computing, Vienna, Austria. ISBN 3-900051-07-0.

Rand DM, Haney RA, Fry AJ (2004) Cytonuclear coevolution: the genomics of cooperation. Trends in Ecology and Evolution 19, 645–653.

Ravinet M, Faria R, RK B, et al. (2017) Interpreting the genomic landscape of speciation: a road map for finding barriers to gene flow. Journal of evolutionary biology.

Riley LG, Cooper S, Hickey P, et al. (2010) Mutation of the mitochondrial tyrosyl-tRNA synthetase gene, YARS2, causes myopathy, lactic acidosis, and sideroblastic anemia—MLASA syndrome. The American Journal of Human Genetics 87, 52–59.

Roux C, Fraïsse C, Romiguier J, et al. (2016) Shedding Light on the Grey Zone of Speciation along a Continuum of Genomic Divergence. PLoS biology 14, e2000234.

Sambatti J, Ortiz Barrientos D, Baack EJ, Rieseberg LH (2008) Ecological selection maintains cytonuclear incompatibilities in hybridizing sunflowers. Ecology letters 11, 1082–1091.

Schrider D, Shanku AG, Kern AD (2016) Effects of linked selective sweeps on demographic inference and model selection. bioRxiv, 047019.

Schwander T, Libbrecht R, Keller L (2014) Supergenes and Complex Phenotypes. Current Biology 24, R288–R294.

Seehausen O, Butlin RK, Keller I, et al. (2014) Genomics and the origin of species. Nature Review Genetics 15, 176–192.

Singhal S, Leffler EM, Sannareddy K, et al. (2015) Stable recombination hotspots in birds. Science 350, 928–932.

Sloan DB, Fields PD, Havird JC (2015) Mitonuclear linkage disequilibrium in human populations. Proceedings of the Royal Society B-Biological Sciences 282, 20151704.

Sloan DB, Havird JC, Sharbrough J (2016) The On Again, Off Again Relationship between Mitochondrial Genomes and Species Boundaries. Molecular Ecology.

Smith J, Coop G, Stephens M, Novembre J (2016) Estimating time to the common ancestor for a beneficial allele. *bioRxiv*, 071241.

Soria-Carrasco V, Gompert Z, Comeault AA, et al. (2014) Stick insect genomes reveal natural selection’s role in parallel speciation. Science 344, 738–742.

Stier A, Bize P, Roussel D, et al. (2014) Mitochondrial uncoupling as a regulator of life-history trajectories in birds: an experimental study in the zebra finch. The Journal of experimental biology 217, 3579–3589.

Sunnucks P, Morales HE, Lamb AM, Pavlova A, Greening C (2017) Integrative Approaches for Studying Mitochondrial and Nuclear Genome Co-evolution in Oxidative Phosphorylation. Frontiers in Genetics 8, 25.

Toews DP, Mandic M, Richards JG, Irwin DE (2014) Migration, mitochondria, and the yellow-rumped warbler. Evolution 68, 241–255.

Villemereuil P, Gaggiotti OE (2015) A new FST based method to uncover local adaptation using environmental variables. Methods in Ecology and Evolution 6, 1248–1258.

Wang J (2017) The computer program STRUCTURE for assigning individuals to populations: easy to use but easier to misuse. Molecular ecology resources.

Warren WC, Clayton DF, Ellegren H, et al. (2010) The genome of a songbird. Nature 464, 757–762.

Weir BS, Cockerham CC (1984) Estimating F-statistics for the analysis of population-structure. Evolution 38, 1358–1370.

Wolf JB, Ellegren H (2016) Making sense of genomic islands of differentiation in light of speciation. Nature Reviews Genetics.

Wolff JN, Ladoukakis ED, Enríquez JA, Dowling DK (2014) Mitonuclear interactions: evolutionary consequences over multiple biological scales. Philosophical Transactions of the Royal Society B: Biological Sciences 369, 20130443.

Wu CI (2001) The genic view of the process of speciation. Journal of evolutionary biology 14, 851–865.

Yeaman S (2013) Genomic rearrangements and the evolution of clusters of locally adaptive loci. Proceedings of the National Academy of Sciences 110, E1743–E1751.

Yip CY, Harbour ME, Jayawardena K, Fearnley IM, Sazanov LA (2011) Evolution of respiratory complex I: “supernumerary” subunits are present in the alpha-proteobacterial enzyme. J Biol Chem 286, 5023–5033.

Zhu J, Vinothkumar KR, Hirst J (2016) Structure of mammalian respiratory complex I. Nature 536, 354–358.

